# Defensive symbiosis and the evolution of virulence

**DOI:** 10.1101/422436

**Authors:** Paul G. Nelson, Georgiana May

## Abstract

Although environments rife with enemies should cause selection for defensive traits, such enemy-rich environments should also select for greater virulence in co-occurring symbionts, yet many defensive symbionts cause little to no damage while protecting their hosts from enemies. Thus, co-infection of a defensive symbiont and a parasite is predicted to select both for increased virulence in co-infecting symbionts and for increased defense in the protective symbiont. Why then do we observe defensive mutualists that protect hosts while causing little damage? To address this question, we build a symbiont-centered model that incorporates the evolution of two independent traits: defense and virulence. Virulence is modeled as a continuous trait spanning mutualism (negative virulence) and parasitism (virulence) and thus accounts for the entire range of direct effects that symbionts have on host mortality. Defense is modeled as a continuous trait that ameliorates the costs to the host associated with co-infection with a deleterious parasite. We obtain the counterintuitive result that the evolution of increased defense in one symbiont largely leads to the evolution of lower virulence in both symbionts and may even facilitate pathogens evolving to mutualism. However, results are context-dependent and when defensive traits are costly, the evolution of greater defense may also lead to higher virulence.

## Body

The recent explosion of research in the “microbiome” makes apparent that hosts rarely, if ever, exclusively interact with a single symbiont, and that multiple infections are the rule rather than the exception (Afkhami et al. 2014; Arnold et al. 2003; Bordenstein and Theis 2015; Saikkonen 2007). Symbionts that defend their host are commonly observed across the microbiomes plant and animal systems (Balmer et al. 2009; Betts et al. 2016; Carroll 1988; Gerardo and Parker 2014; Jaenike et al. 2010; Madden and Young 1992; Vittecoq et al. 2012). (Arnold et al. 2003; Clay and Schardl 2002; Oliver et al. 2008). While defensive symbioses are both common and of increasing importance to disease control and host health (Ford and King 2016), we know too little of the evolutionary and ecological consequences of symbiont interactions within a host to predict outcomes for virulence and the maintenance of defensive symbioses (May and Nelson 2014). To understand the impact of interactions among microbial symbionts on the evolution of virulence and defense, we lay out a model that includes two different processes: the direct effects of the symbiont on the host, incorporated as virulence, and the indirect effects of a symbiont killing or inhibiting co-infecting symbionts, incorporated as defense of the host.

Expectations for the evolution of a symbiont’s impact on host health, virulence, are predicated primarily on theory developed for host-parasite interactions and assume a trade-off between transmission rate and infection duration (Alizon et al. 2009; Anderson and May 1978; Ewald 1987). Further, under co-infection, competition among parasites within hosts or for hosts in a population should cause selection for increased parasite virulence (Alizon et al. 2009; Barrett et al. 2011). Generally, symbionts that harm their host to gain reproduction are expected to evolve to intermediate levels of virulence at equilibrium because of the trade-off between symbiont transmission rate and host mortality (Alizon et al. 2009; Barrett et al. 2011). While trade-offs have received far less attention for beneficial symbionts, the classic tradeoff applies if we treat mutualism as negative virulence and assume that a mutualist’s transmission decreases with the degree to which that mutualist aids its host (Friesen and Mathias 2010; Nelson and May 2017; Orivel et al. 2017). Still, because theory for the evolution of mutualism and parasitism often treat these extremes of the possible range of virulence separately, the evolution of virulence and defense in a community context of the microbiome remains problematic.

To understand the evolution of symbiont virulence along the continuum from parasitism to mutualism, we developed the concept of shared costs (Nelson and May 2017) as a common “currency” accounting for the costs and benefits of multiple symbionts within a shared host. For example, as parasites sequester resources (Choisy and de Roode 2010) and decrease host quality (Ebert and Mangin 1997), the costs of damage to the host are shared among all co-occurring symbionts in that host (Nelson and May 2017). A large body of theoretical and empirical results demonstrate that infection by multiple parasites should cause selection for increased pathogen virulence when the costs of virulence are shared and each symbiont receives the individual benefits of its own virulence (Alizon et al. 2009; Bremermann and Pickering 1983; Ebert and Mangin 1997; Friesen and Mathias 2010; Mosquera and Adler 1998; Nelson and May 2017; van Baalen and Sabelis 1995; West et al. 2002). Building on results of Nelson and May (2017), we study the impacts of interactions between defensive and parasitic symbionts on the evolution of virulence and defense, mediated through shared costs of virulence.

Considering both the predictions from models addressing the evolution of defense and those model predictions for evolution of virulence produces a conundrum. On one hand, we expect diverse communities of interacting symbionts to promote the evolution of defense (Kerr et al. 2002; Kirkup and Riley 2004). On the other hand, competition among interacting symbionts generate the same conditions under which more virulent symbionts are expected to evolve (Frank 1996). Models regarding the evolution of defense predict that co-infection of the same or different symbiont species will cause selection for defensive mutualists (Fenton et al. 2011; Jones et al. 2007), while models regarding the evolution of virulence predict the evolution of virulent pathogens (Mosquera and Adler 1998) under the same conditions. The possibility of defensive pathogens has been neglected until recently, with results of an experimental co-evolutionary study demonstrating that pathogenic symbionts may indeed evolve traits to suppress other symbionts growth (Ford et al. 2016).

To address the gap between the predictions from theory for the evolution of defensive traits and those for evolution of virulence, we build a model that incorporates defense, virulence, and co-evolution between symbiont species under infection by multiple symbionts of potentially different species. We incorporate virulence as a continuum between parasitism (positive virulence) and mutualism (negative virulence) as in (Nelson and May 2017), and extend that model to include symbiont defense of the host. (Lipsitch et al. 1996). In our model, we make no explicit assumptions regarding vertical or horizontal transmission; we merely track symbiont birth and death rates and thus take a different approach than previous work (Jones et al. 2011; Lively et al. 2005; Sorrell et al. 2009), which analyzed vertically transmitted defensive symbionts. Finally, we use the concept of the shared costs of virulence (Nelson and May 2017) to model the impacts of multiple infections, both of the same and different symbiotic species, on the evolution of virulence and defense traits of symbionts. By using a continuous scale for virulence and defense, we aim to capture the highly varied interactions between symbionts observed in nature. We develop a model for the evolution of virulence and defense in the context of a microbial community to ask if evolvable defense necessarily leads to mutualism.

## Methods

**Table 1:**
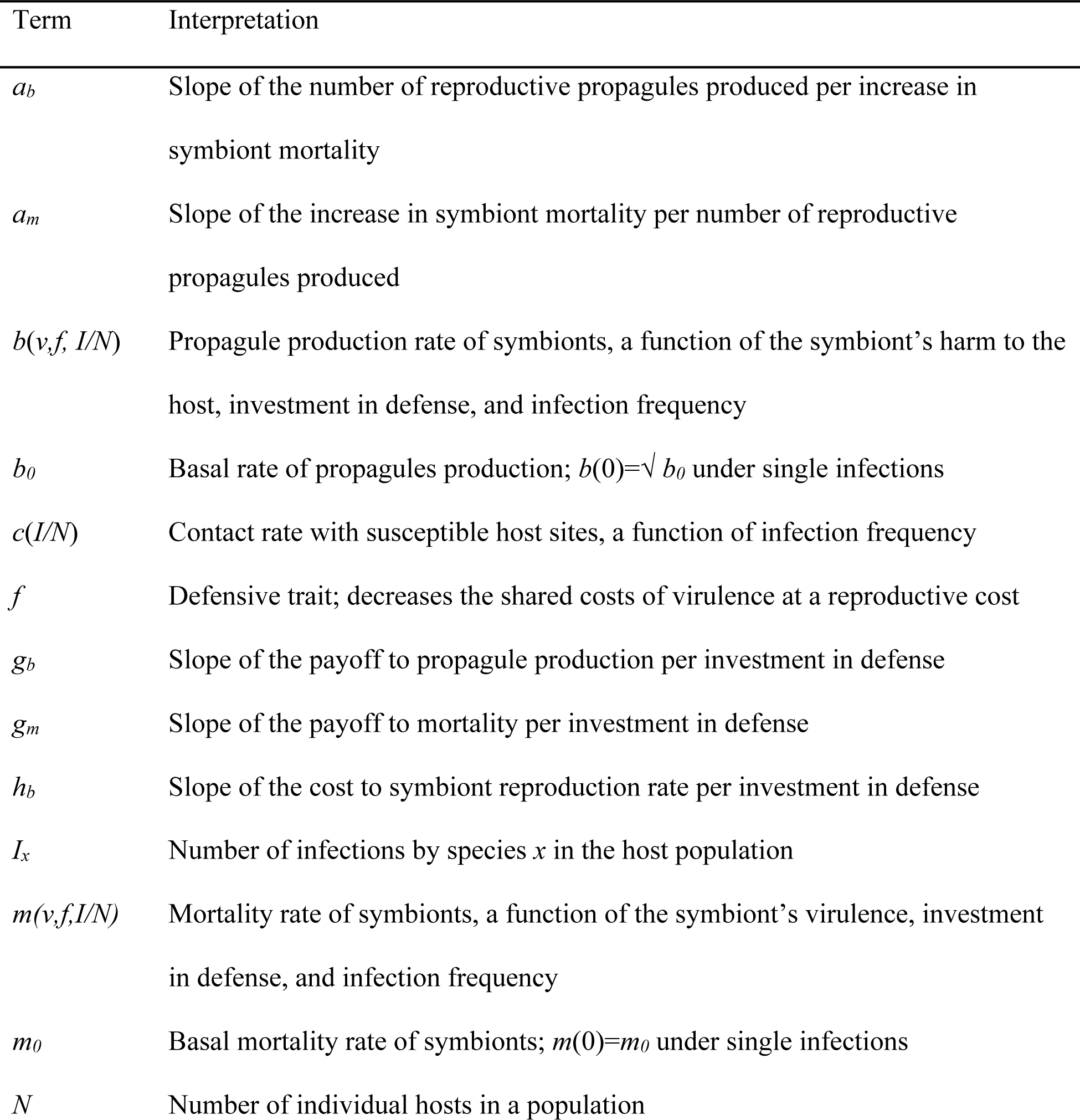
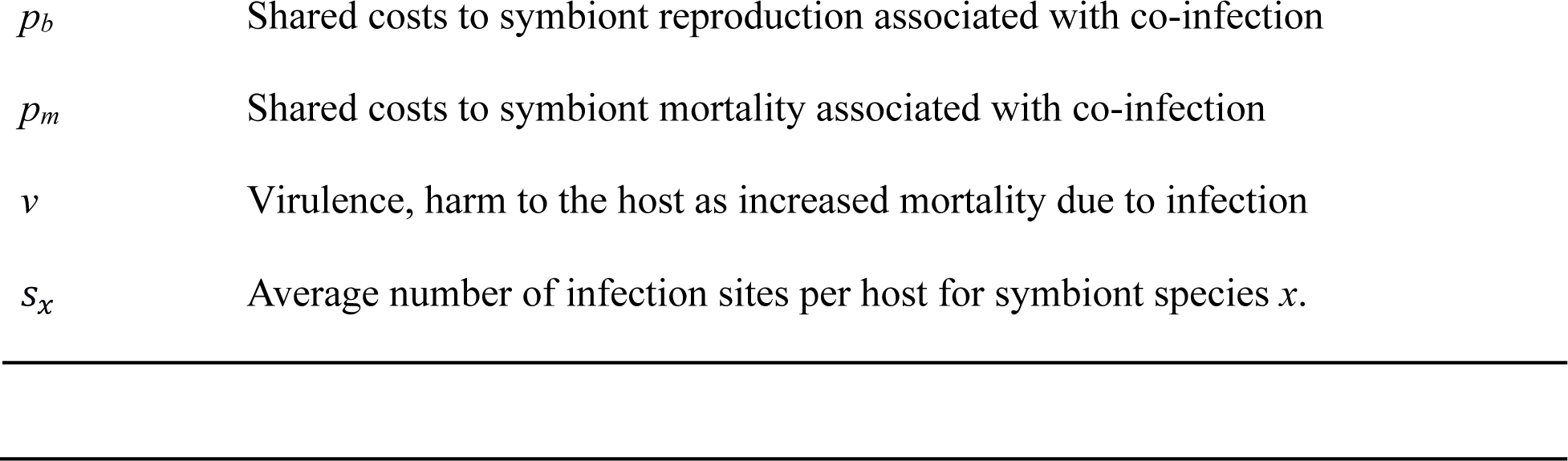
Terms used

Following Nelson and May (2017), we lay out an approach to symbiont evolution that uses host population density to feedback on virulence and transmission (Lenski and May 1994) but here, focuses on the impacts of interactions among symbionts in driving the costs of virulence. We then incorporate expressions for the impacts of evolvable defense into specific functions for symbiont birth and mortality for numerical analysis of stable states at differing levels of costs and defense. Infections within hosts of strain *i* of species *x* “die” at rate *m* and produce infectious propagules at rate *b,* per unit time. Propagules transmit between hosts and establish new infections at rate *c*, making *bc* the effective symbiont transmission rate. Together, the functions *b c*, and *m* yield the expected change in number of infections for each strain *i* of symbiont species *x* (*I*_*ix*_):

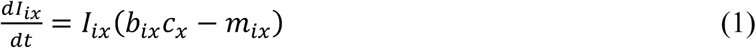

Assuming the probability of establishing a new infection is density dependent,

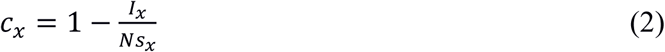

where *s*_*x*_ determines the density dependence of the probability of establishing new infections, and *Ns*_*x*_ is analogous to the carrying capacity in the logistic growth function (Nelson and May 2017). Equations 1 and 2 produce a negative density dependent feedback similar to that in (Lenski and May 1994) and resulting in an ecological equilibrium (*dI*/*dt* = 0) where:

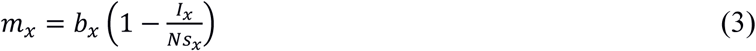

Next, we develop specific functions for *b* and *m* that incorporate both virulence (*v*) and defense (*f*), and then use these to solve for equilibria numerically. Both symbiont reproduction and mortality are increasing functions of virulence *v* and defense *f* such that *m* increases faster than *b* as either *v* or *f* increase (*dm*/*dv*>*db*/*dv* and *dm*/*df*>*db*/*df*), so that the per capita change in infection frequency (*dI/Idt*) is a convex function of *v* and *f*. We develop our functions in three steps: first we examine the model under single infections, then multiple infections by adding the shared costs of virulence, and finally we incorporate symbiont defense of the host.

Under single infections, mortality is a linear function of virulence, modulated by the coefficient *a*_*mx*_, and *m*_*0*_, the basal symbiont mortality rate. Reproduction is the square root of virulence, again modulated by a coefficient *a*_*bx*_, and basal symbiont propagule production rate *b*_*0*_. The basal rate of propagule production, *b*_*0*_, acknowledges that symbionts may reproduce without net harm to the host (zero virulence), and allows the model to encompass evolution of the full range of symbiont’s effects on the host, from mutualism (negative virulence) to parasitism (positive virulence) (Nelson and May 2017). Under single infections:

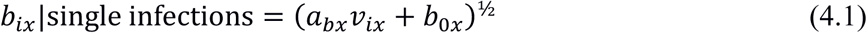

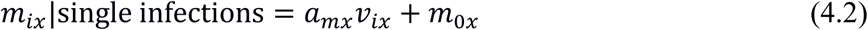

Next, we incorporate the impacts of multiple symbionts co-infecting a host using a term for the shared costs of virulence (*p*) that allows co-evolution between symbiont species. The shared costs of co-infection may affect symbiont birth (*p*_*b*_) or mortality (*p*_*m*_). If both *p*_*b*_ and *p*_*m*_ are zero, symbionts have no effect on each other, and the model results will be the same as a single-infection system. If *p*_*b*_ is positive, co-infecting symbionts compete for host resources and affect each other’s reproduction. If *p*_*m*_ is positive, symbionts indirectly affect each other by affecting host mortality, thus affecting the mortality of other symbionts in the host. To obtain the total shared costs of virulence, we multiply the coefficients *p*_*b*_ and *p*_*m*_ by the average virulence of symbionts infecting a host population. Starting with the mean virulence of symbiont species k, 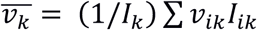, we take the sum of virulence across species, weighted by the infection frequency of that species 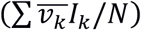, yielding the total shared costs 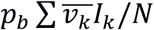 and 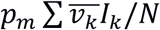. It is important to note that we are assuming, for simplicity, an even distribution of symbionts across the host population.

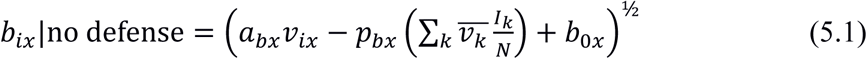

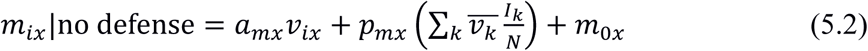

Finally, we incorporate defense (*f*) with both costs and benefits to the symbiont. The cost of defense to symbiont reproduction is determined by the parameter *h*_*b*_ > 0. Defense of the host benefits the defending symbiont by decreasing the shared costs associated with a co-infecting deleterious parasite. Because we are not focusing on specific mechanisms of defense, we use a relatively simple function in which an increased investment in *f* decreases the shared costs of virulence without affecting other terms. While it is mathematically possible for defense to be negative and thus model facilitation, here we focus on host defense by limiting the analysis to *f* ≥ 0. We divide *p*_*b*_ and *p*_*m*_ by investment in the defensive trait, multiplied by a coefficient *g*_*b*_ and *g*_*m*_, plus one. If both *g*_*b*_ and *g*_*m*_ are zero, investment in defense yields no benefit and defense is not evolvable. Incorporating defense yields:

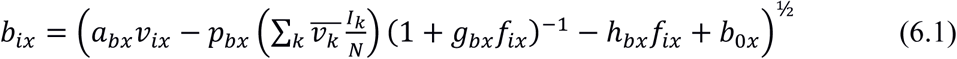

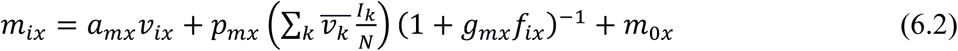

Note that in equations 2, 6.1, and 6.2, total host population size *N* appears only in the context of infection frequency *I*/*N*, meaning the results are independent of host population size *per se*. Therefore, we need not assume that host population size is constant, only that there are no forces affecting the infection frequency that are not accounted for by the parameters given here.

### Evolutionary stable states

Evolutionary equilibrium occur when the derivative of the per capita change in infection frequency with respect to *v* and *f* is zero (Abrams et al. 1993). An equilibria is an evolutionary stable state when it is at a fitness maximum, as indicated by a definite negative Hessian matrix (Debarre et al. 2014). First, we examine the first partial derivatives of *dI*_*ix*_/*Idt* for equilibria. To simplify the expression of equations below, we represent total shared costs with 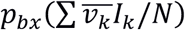 and 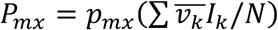:

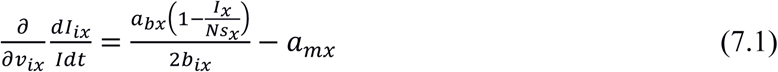

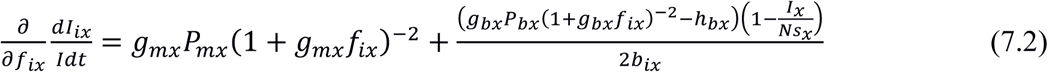

Equation 7.1 is zero when 2*b*_*ix*_*a*_*mx*_ = *a*_*bx*_(1 – *s*_*x*_*I*_*x*_/*N*). Equation 7.2 shows that when total shared costs are negative (*P*_*mx*_ and *P*_*bx*_≤ 0) due to co-infection with mutualists, then ∂/∂*f*_*ix*_(*dI*_*ix*_/*Idt*) < 0. In this case, selection always favors lower defense, defense has no evolutionary equilibrium and investment in defense tends toward zero.

Next, we determine if the equilibrium in *dI*_*ix*_/*Idt* with respect to virulence and defense is at a minimum or a maximum. As shown in Debarre et al. (2014), a Hessian matrix uses the second derivatives of a fitness function, here *dI*_*ix*_/*Idt*, to determine if a stable state sits at a fitness peak, a fitness minimum, or at a saddle. An evolutionary maximum occurs when the Hessian matrix is definite negative. A sufficient condition for a definite negative a Hessian matrix is that the second derivatives by *v* and *f* are negative, and ∂^2^/∂*f*_*ix*_∂*v*_*ix*_(*dI*_*ix*_/*Idt*) = ∂^2^/∂*v*_*ix*_∂*f*_*ix*_(*dI*_*ix*_/*Idt*) = 0. To determine if the Hessian matrix is definite negative, we examine the second derivatives of *dI*_*ix*_/*Idt*:

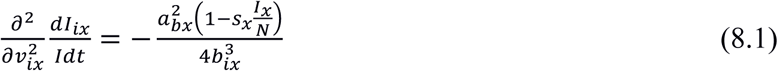

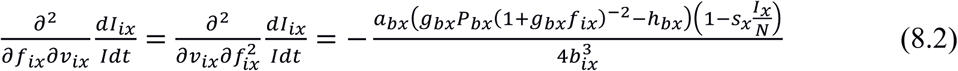

Under the equilibrium condition where ∂/∂*f*_*ix*_(*dI*_*ix*_/*Idt*) = 0 (Eq. 7.2), Equation 8.2 can be rewritten as:

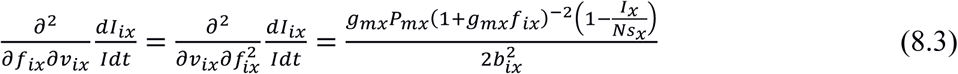

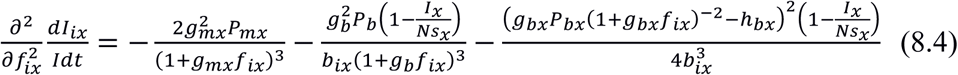

Equation 8.1 is negative under all conditions. Equation 8.2 is zero when *P*_*bx*_ = 0 and Equation 8.3 is zero when *P*_*mx*_ = 0, meaning ∂^2^/∂*f*_*ix*_∂*v*_*ix*_(*dI*_*ix*_/*Idt*) = ∂^2^/∂*v*_*ix*_∂*f*_*ix*_(*dI*_*ix*_/*Idt*) = 0 when either *p*_*bx*_ or *p*_*mx*_ is zero. Lastly, Equation 8.4 is negative when either *P*_*b*_ or *P*_*m*_ are positive. Therefore, when total shared cost are positive and affect either *b* or *m* (either *P*_*mx*_>0, or *P*_*bx*_>0, but not both), the Hessian matrix of *dI*_*ix*_/*Idt* is definite negative, indicating that stable states defined by the roots of equations 7.1 and 7.2 are fitness peaks, and evolutionary stable states (Debarre et al. 2014).

### Solving for equilibria

As closed-form solutions can only be expressed under single infections, we used a slightly modified version of Broyden’s method (Broyden 1965) to find evolutionary stable states at ecological equilibrium. Broyden’s method is a quasi-Newton search algorithm for finding maxima of a function in multidimensional space, which uses the slope of secants to approximate the Jacobian matrix used in Newton’s method (Press et al. 2007). At each step *n* of the search, *v*_*x*_(*n* + 1) = *v*_*x*_(*n*) + *β*(*dI*_*x*_/*I*_*x*_*dt*(*v*_*x*_(*n*) + *ε*) - *dI*_*x*_/*I*_*x*_*dt*(*v*_*x*_(*n*) - *ε*))/*ε*, where *ε* is a small number (*ε*=0.0001 here) that determines the accuracy of the search and *β* determines the speed of the search (*β*=0.0002 here). Likewise, *f*_*x*_(*n* + 1) = *f*_*x*_(*n*) + *β*(*dI*_*x*_/*I*_*x*_*dt*(*f*_*x*_(*n*) + *ε*) - *dI*_*x*_/*I*_*x*_*dt*(*f*_*x*_(*n*) - *ε*))/*ε*. Instead of using the secant to increment *I*_*x*_, we use *dI/dt*: *I*_*x*_(*n* + 1) = *I*_*x*_(*n*) + *βdI*_*x*_/*dt*. The search is halted when *dI*_*x*_/*I*_*x*_*dt* is lower for all new values of *v*_*x*_ and *f*_*x*_ tested, *dI*_*x*_/*dt* = 0, and *dI*_*x*_/*I*_*x*_*dt*(*v*_*x*_ ± *ε*) < *dI*_*x*_/*I*_*x*_*dt*(*v*_*x*_), *dI*_*x*_/*I*_*x*_*dt*(*f*_*x*_ ± *ε*) < *dI*_*x*_/*I*_*x*_*dt*(*f*_*x*_). A limitation of using Broyden’s method on this model is that if *v* is negative (mutualism), the results of the search algorithm can diverge from analytical predictions above. Specifically, if total shared costs, 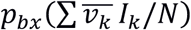 or 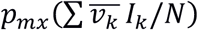, are negative, which only occurs when *v* is negative, Equation 7.2 predicts that *f* will decrease to zero. However, if *v* is negative, *m c,* and *dI/Idt* can become zero, which makes the slopes of all secants in the local vicinity zero, and the search algorithm is insensitive to any further changes in *v* or *f* even if such changes would increase *b*. In effect, the search becomes “stuck” at the values of *v* and *f* that first result in *m*=0. To prevent the search from becoming stuck, we floor the possible values of *c* at a number close to, but not equal to zero when calculating the secants (0.0005 here), allowing the algorithm to maximize *b* even if *m*=0.

For the results presented below, we examine the evolution of virulence and defense in two species. One symbiont species has a positive basal rate of propagule production (*b*_*0*_ >0) and thus has the potential to evolve mutualism and cause decreased host mortality rates. The second species has zero basal reproduction rate (*b*_*0*_=0) and thus can only be an “obligate” pathogen that gains reproduction at the cost of increased host mortality. Initial search values (*v*_*x*_=0.1, *I*_*x*_*/N*=0.1, *f*_*x*_=0) were chosen to correspond to pathogenic symbionts with no investment in defense that occur initially at a low infection frequency. We initialize the search with pathogenic symbionts to provide a conservative view on conditions for the evolution of mutualism. We limit our analysis to scenarios where either, but not both, *p*_*bx*_ or *p*_*mx*_ is zero, because the Hessian matrix is not definite negative when both *p*_*bx*_ and *p*_*mx*_ are positive, meaning evolutionary stable states may not exist under those conditions. For all results we use *a*_*b*_=4, *a*_*m*_=1, *m*_*0*_=0.1, and *h*_*b*_=0.1; values of *b*_*0*_, *p*_*b*_, *p*_*m*_, *g*_*b*_, and *g*_*m*_ are specified in the figure legends.

## Results

We examined equilibrium states for virulence and defense in an obligate pathogen (*b*_*0*_=0) and a potential mutualist (*b*_*0*_ =0.3) symbiont species. First, we assessed outcomes if each of these symbiont species is evolving alone, in which case shared costs affect only members of the same species within a host. Then, we examined a scenario in which the pathogen and potential mutualist co-evolve, in which case shared costs from one species affect the other species. We contrast results for the evolution of virulence when symbionts can also evolve defense to the results obtained when defense cannot evolve (*f*=0, *g*_*b*_=*g*_*m*_=0). For a thorough investigation of the effect of shared costs of virulence without defense, see Nelson and May (2017).

### Shared costs within species

We first examine the evolution of virulence and defense when each species evolves separately. Here, each species if affected by the shared costs of only that species (e.g. for two species, *a* and *b*, shared costs are 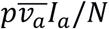 for species *a* and 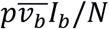 for species *b*). Overall, Fig. 1 shows that patterns of evolved virulence (A, B) and infection frequency (C, D) are similar whether shared costs affect symbiont reproduction (*p*_*b*_; left column) or symbiont mortality (*p*_*m*_; right column). As we might expect, a potential mutualist (*b*_*0*_ *>* 0; black lines) and an obligate parasite (*b*_*0*_ *=* 0; grey lines) have different evolutionary responses to increasing shared costs and evolvable defense. When shared costs are low, the potential mutualist (black line Fig. 1 A, B) evolves negative virulence. As shared costs continue to increase, mutualism is lost, and the symbiont evolves positive virulence or parasitism. Similarly, the pathogen (dashed grey line Fig. 1 A, B) evolves greater virulence with increased shared costs, but unlike the mutualist, never evolves negative virulence. In both the pathogen and the potential mutualist, the ability to evolve defense partially offsets the costs of virulence and lower levels of virulence evolve even under high shared costs. The added flexibility of modeling the evolution of virulence and defense as independent traits allows us to show that parasitic symbionts may evolve defense traits that reduce host harm caused by other virulent symbionts. For the obligate pathogen, higher shared costs to symbiont reproduction or mortality always lead to lower infection frequencies (dashed grey line Fig. 1 C, D). For the potential mutualist, in contrast, low to moderate shared costs to this symbiont’s reproduction or mortality actually lead to increased infection frequencies (dashed black line Fig. 1 C, D), because the mutualist individuals decrease host mortality, increase each other’s reproduction within the host, and thus increase transmission rates. The evolution of defensive traits lowers the impact of shared costs, and thus evolvable defense (solid lines Fig. 1 C, D) results in greater infection frequencies for both parasites and mutualists than would otherwise be achieved without defense.

**Figure 1:**
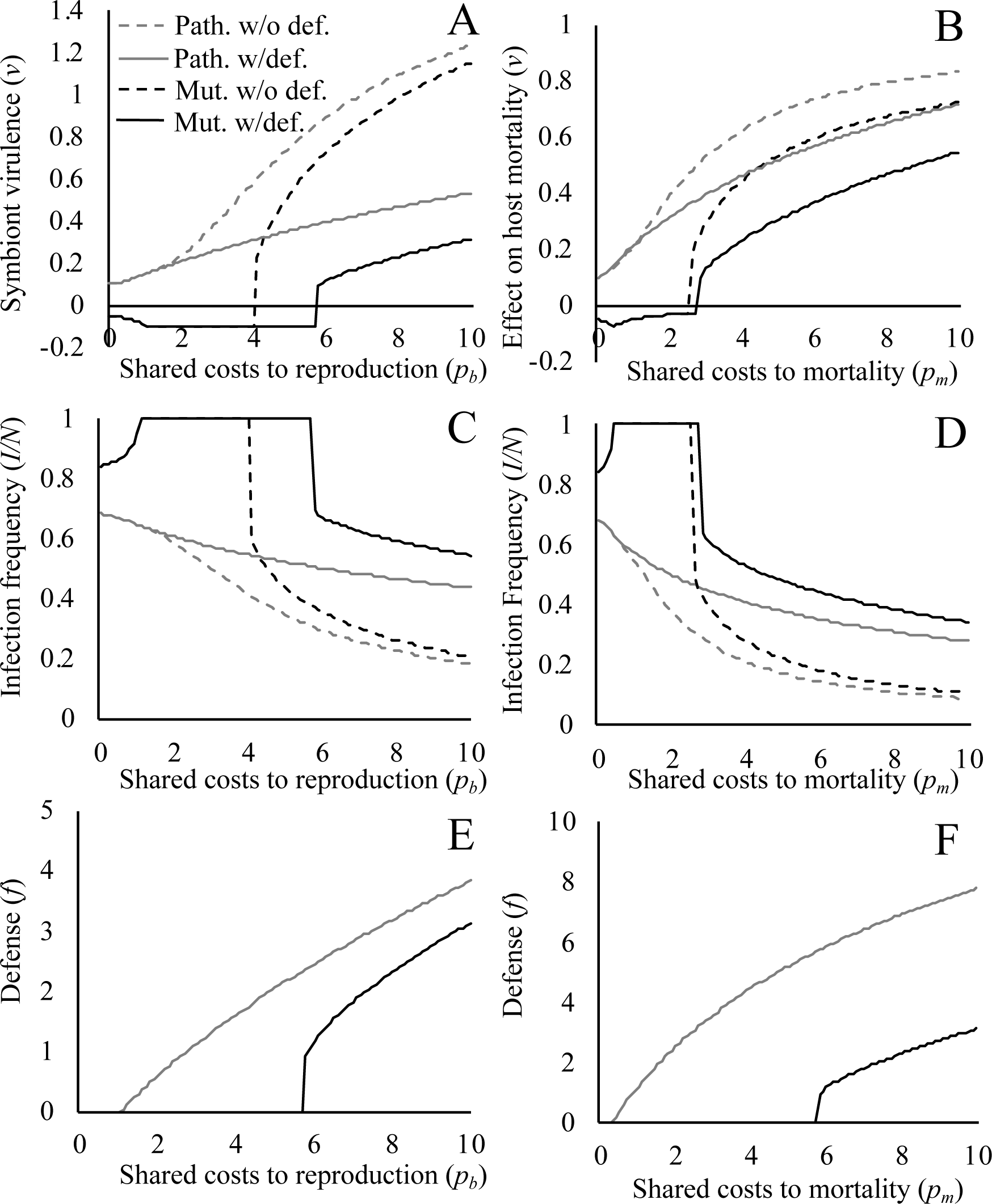
Evolutionary stable states for each species evolving alone. The costs of virulence are shared within species for either an obligate pathogen (grey, *b*_*0*_=0) or a potential mutualist (black, *b*_*0*_=0.3). Evolutionary stable states with evolvable defense (*g*_*m*_=1 or *g*_*b*_=1) are shown with solid lines, evolutionary stable states without evolvable defense (*f*=0, *g*_*m*_=0 and *g*_*b*_=0) are shown with dashed lines. The results for symbiont virulence (A and B), infection frequency (C and D), and evolved defense (E and F) at equilibrium are shown for increasing values of shared costs to symbiont reproduction (*p*_*b*_; A, C, E) and to symbiont mortality (*p*_*m*_; B, D, F).

Evolvable defense tempers the effects of shared costs for both the potential mutualist and the pathogen. Pathogenic species (grey line) invest in defense at relatively low levels of shared costs, while shared costs must be large before the potential mutualist (black line) invests in defense (Fig. 1 E, F). Similar results are obtained whether shared costs effect symbiont reproduction or mortality. The potential mutualist only evolves defense after mutualism is lost, an unsurprising result considering co-infecting with a fellow mutualist is beneficial and defending against that mutualist would be counterproductive. Interestingly, the potential mutualist (Fig. 1 A, B, black lines) maintains negative virulence at higher shared costs with defense than without defense, even though investment at equilibrium in defense is zero. As shared costs continue to increase, defense allows the evolution of less damaging parasites even after the symbiont switches to a pathogenic strategy (*v*>0). While we observe that defense most often leads to the evolution of lower virulence, we do note that when shared costs to symbiont mortality are low (*p*_*m*_ < 1), evolvable defense actually leads to slightly more damaging pathogens (Fig. 1 B, grey lines). This unexpected result is explored further below.

### Coevolution between symbiont species

We next examine the effect of shared costs on co-evolution between two species, one a potential mutualist and the other an obligate pathogen. To isolate the effects of interactions between species from the effects of interactions within species, each species is affected by the total shared costs of virulence of the other species. Specifically, for two species, *a* and *b*, shared costs for species *a* are 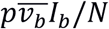 and for species *b*, 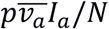.

When costs are shared between species, the presence of a mutualist decreases the mortality rate, or increases the propagule production rate, of the pathogen, leading to the evolution of decreased virulence in the obligate pathogen (Fig. 2 A, B). At low to moderate levels of shared costs, the presence of a mutualist species facilities the evolution of lower, and even negative, virulence (mutualism) in the erstwhile pathogen. In this case, the mutualist causes an increase in the co-infecting pathogen’s reproduction, effectively making the basal propagule production rate >0 for the pathogen and allowing it to evolve negative virulence. When shared costs are high, the potential mutualist evolves positive virulence, and both species become parasites. While the results are similar whether costs are attributed to symbiont reproduction (*p*_*b*_) or mortality (*p*_*m*_), it is interesting that when shared costs between two species affect symbiont mortality (Fig. 2 B), the presence of a mutualist may not only lead to the evolution of decreased virulence in the pathogen but under some conditions (*p*_*m*_ ∼ 1), may also lead to commensalism (*v* ∼ 0) in the pathogen. However, in contrast to when shared costs affect symbiont reproduction (Fig. 2 A), the pathogen never evolves mutualism when shared costs affect mortality

**Figure 2:**
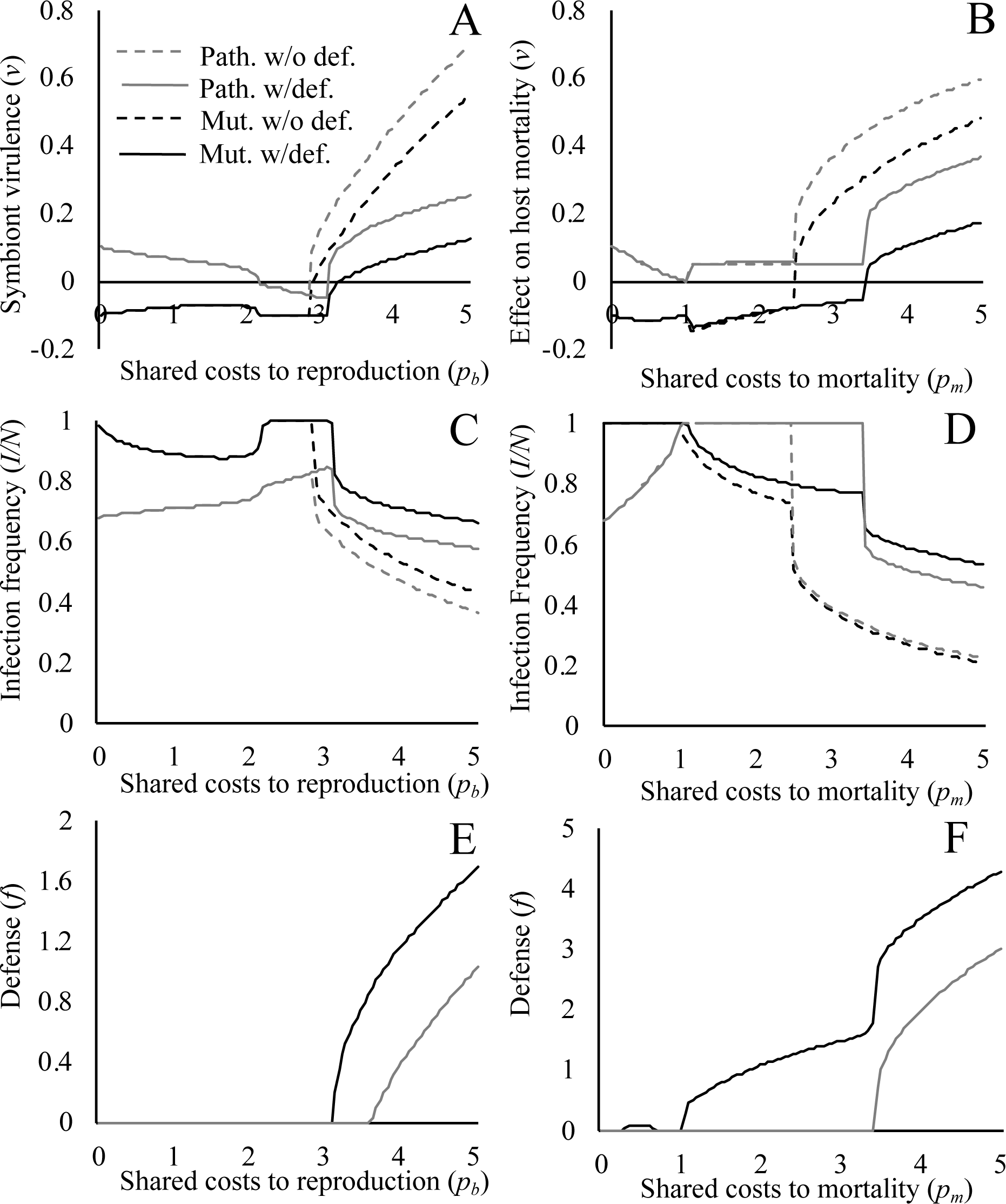
Evolutionary stable states when two species co-evolve. The evolved virulence (A and B), infection frequency (C and D), and evolved defense (E and F) at equilibrium are shown for increasing values of shared costs to reproduction (*p*_*b*_; A, C, E) and mortality (*p*_*m*_; B, D, F). The costs of co-infection are shared between a population of obligate pathogens (grey, *b*_*0*_=0) and a population of potential mutualists (black, *b*_*0*_=0.3) Evolutionary stable states with evolvable defense (*g*_*b*_=1, left column or *g*_*m*_=1, right column) are shown with solid lines and the ESS without evolvable defense (*f*=0, *g*_*b*_=0 and *g*_*m*_=0) are shown with dashed lines.

Shared costs also affect infection rates of pathogens and mutualists differently (Fig. 2 C, D). Under low to moderate shared costs affecting symbiont reproduction, as shared costs increase, the infection frequencies (*I/N*) for the pathogen increase, and infection frequencies for the mutualist decrease. When shared costs are high (*p*>3), both species suffer decreasing infection frequencies as shared costs increase further and both species evolve greater virulence (Fig. 2 C). When shared costs affect mortality (Fig. 2 D), co-infection with a mutualist leads to increasing infection frequencies in the pathogen only when shared costs are low. As shared costs increase, the deleterious effects of the pathogen lead to decreasing infection frequencies in the mutualist. Finally, when shared costs are high, infection frequencies in both species decrease.

When shared costs between species affecting symbiont reproduction are low, neither the mutualist nor the pathogen evolve defense (Fig. 2 E) and evolvable defense has little to no effect on the evolution of virulence. As shared costs increase, an investment in defense by the mutualist mitigates the costs of co-infecting with the pathogen and allows the potential mutualist to evolve negative virulence at higher levels of shared costs than when defense is not evolvable (Fig. 2 A, *p*_*b*_≈3.1 vs 2.9, and B, *p*_*m*_≈3.4 vs 2.4). The pathogen only invests in defense when shared costs are high enough to cause the potential mutualist to evolve positive virulence and become a pathogen itself, here evolvable defense results in lower virulence than when defense is not evolvable.

### Effects of evolvable defense on mutualism

The results for shared costs within species above yield two unexpected outcomes. First, evolvable defense facilitates the evolution of mutualism even though investment in defense at equilibrium is zero (compare Fig. 1 A and E for the region 4<*p*_*b*_<5.8, black lines). To understand the process by which defense might facilitate the evolution of mutualisms but come to zero at equilibrium, Fig. 3 shows the evolved level of virulence and investment in defense over time for a potential mutualist under a single value of shared costs (*p*_*b*_=5). Starting from a pathogenic state with low infection frequency, selection favors more damaging strains. Without evolvable defense, virulence toward the host increases and then plateaus, with the potential mutualist evolving to a pathogenic steady state. In contrast, with evolvable defense, the levels of virulence and of defense peak and then both decrease, but the investment in defense lasts longer before declining. Once the population reaches a stable mutualistic state, defense may evolve to zero. This counterintuitive result stems from dynamics shown in Equation 8.2, which for the parameters used in Fig. 3 (*p*_*m*_=0; *p*_*m*_>0) can be reduced to:

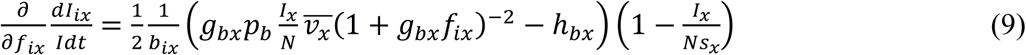

**Figure 3:**
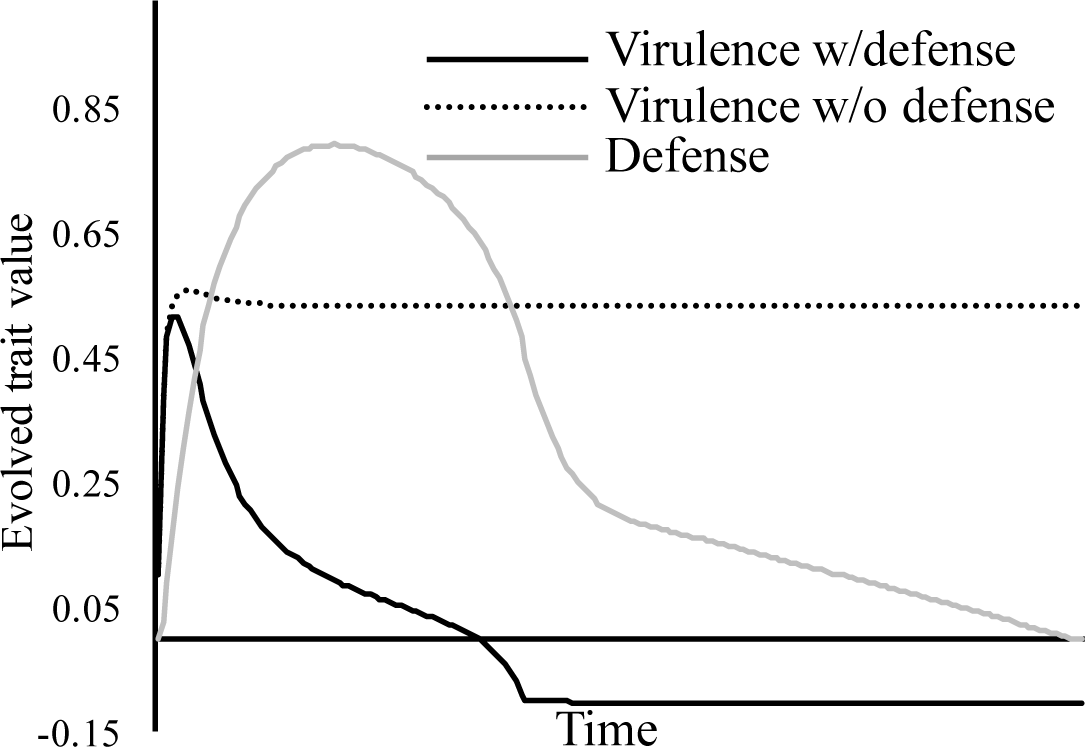
The evolution of mutualism and defense over time with shared costs affecting symbiont reproduction. To investigate how defense can facilitate the evolution of mutualism and then be lost, a time series shows the evolution of virulence in a potential mutualist (*b*_*0*_=0.3; *p*_*b*_=5) with (*g*_*b*_=1, solid line) and without (*f*=0, *g*_*b*_=0, dotted line) evolvable defense. The level of evolved defense (grey) is shown.

As one can see from Equation 9, if 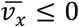, then ∂/∂*f*_*ix*_(*dI*_*ix*_/*Idt*) is negative, indicating that selection favors lower defense. Essentially, the presence of virulent co-infecting symbionts is needed to maintain selection for defensive traits, and once those virulent symbionts are replaced by commensal or mutualist symbionts, selection for defense is relaxed. Thus, defense allows for the evolution of beneficial symbionts, at which point selection purges the defensive trait. This result raises the possibility that transient investment in defense may leave a legacy of mutualism.

Second, when shared costs affect mortality, there is a small region at low shared costs (Fig. 1B; *p*_*m*_ *<* 1) where the parasite evolves slightly greater virulence with evolvable defense than without. While this effect is extremely small, it raises the possibility that under some conditions evolvable defense might lead to more damaging parasites and actually disrupt mutualisms. To determine if defense always facilitates mutualism, we broaden our approach and examine a range of payoffs for investment in defense. Thus far we have examined a single payoff regime where the costs of defense are relatively low compared to the benefits (*h*_*b*_=0.1, *g*_*m*_=1 or *g*_*b*_=1). Because we are primarily interested in the effect of evolvable defense on the evolution of beneficial symbionts, we focus on the critical value of shared costs at which the potential mutualist evolves to a pathogenic state, *p\**. In Fig. 1 A, for example, *p\**≈5.8 with defense and *p\**≈4 without defense. The ratio of *p\** with and without evolvable defense illustrates how the costs of investment in defense affects the evolution of mutualism (Fig. 4). Ratios of *p\** with and without evolvable defense greater than one indicate that the defense trade-off allows mutualism to evolve at higher levels of shared costs, whereas values less than one indicate that evolvable defense disrupts mutualisms and selects for pathogenicity. When shared costs affect reproduction, evolvable defense always preserves mutualisms. However, when shared costs affecting symbiont mortality, when g_m_ is low the defense trade-off can actually disrupt mutualisms and lead to parasitism.

**Figure 4:**
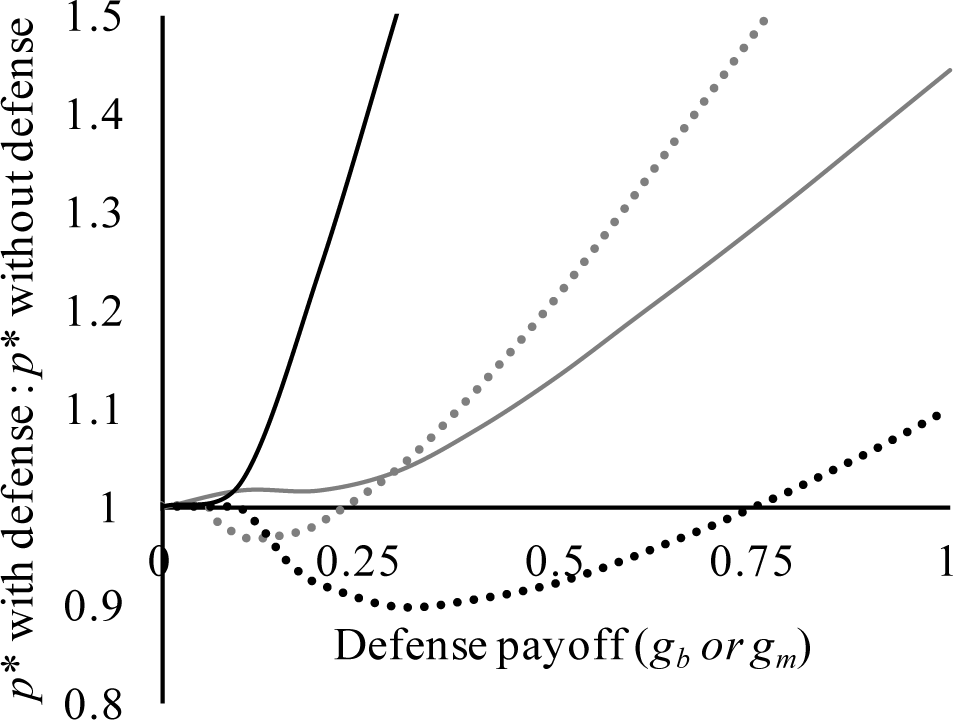
Defense payoff and the evolution of mutualism. To investigate the effect of return on investment in defense (*g*_*m*_ or *g*_*b*_) on the critical value of shared costs at which a potential mutualist evolves parasitism (*p\**), we examine the ratio of *p\** with defense to *p\** without defense. Values above one indicate that evolvable defense facilitates the evolution of mutualism, whereas values below one indicate that evolvable defense hampers the evolution of mutualism. Four scenarios of shared costs are shown: shared costs within a potential mutualist species (black), shared costs between a potential mutualist and an obligate pathogen (grey), shared costs affecting reproduction (solid), and shared costs affecting mortality (dotted).

## Discussion

An important question raised by Lenski and May (1994) is yet unanswered: “Why are some parasites so highly virulent, whereas other parasites, often closely related, are much less harmful to their hosts?” We expand upon the body of literature addressing this question in three ways. Like Jones (2007) and Jones (2011), we expand on Heithaus (1980), Fenton (2011), and Kwiatkowski (2012) by making symbiont virulence and symbiont defense of the host two quantitative traits that evolve together. In our model, the level of protection a symbiont provides its host is variable and subject to selection, allowing us better model the role of natural selection in maintaining defensive mutualisms (Johnson et al. 1997). Additionally, we generalize Jones (2007) and Jones (2011) by incorporating horizontal transmission between hosts. Thus, symbionts in our model can transmit horizontally or vertically and establish multiple infections, with different symbionts of the same or different species.

In order to explain the evolution of low virulence under multiple infections, previous studies have invoked symbiont population structure, specifically relatedness and kin selection (Buckling and Brockhurst 2008; Koskella et al. 2006). Our results speak to an alternative mechanism: evolvable defense of the host can lead to lower virulence and even mutualism. However, we find that evolvable defense can sometimes lead to the evolution of a defensive parasite with greater virulence than it would have without evolvable defense. Such a seemingly counterintuitive result arises when defense is costly, because symbionts compensate for that cost by harvesting more resources from their hosts. Neuhauser and Fargione (2004) obtained similar results by placing symbiotic organisms on a resource gradient to explain observations of symbiotic organisms along a mutualism-parasitism continuum. Most importantly, our results inform the conditions under which lower virulence, mutualisms and commensalism (virulence is 0) are apparent evolutionary stable states. While the focus of microbiome studies has almost exclusively been on beneficial mutualisms, commensalism is by far the most common state and may constitute a vast reservoir of genetic variation in symbiotic populations (May 2016).

When symbiont species co-evolve, our results speak directly to the co-evolution of defensive mutualists in the presence of parasites. Similar to the conclusions of Fenton et al. (2011) for the conditions under which the *Wolbachia* parasite might successfully invade new insect hosts, our results show that if the cost of investment in defense is low and the shared costs of virulence are not too high, both parasitic and mutualistic symbionts may evolve lower virulence. Further, the experimental results of Ford et al. (2016) provide empirical validation of our results as they demonstrate the evolution of lower virulence with defensive symbionts.

### Future directions

In examining these trade-offs we have made a number of simplifying assumptions. Chiefly, we have neglected modeling the role of new mutations affecting virulence and defense. We have also neglected demographic processes such as genetic drift, or the impact of environmental disturbances, either of which may dominate when selection pressures are low or populations are small or spatially structured (Nahum et al. 2011; Prado and Kerr 2008; Verbruggen et al. 2012). Similar models addressing evolution of cooperation have shown that, given sufficient genetic variation, a species may evolve to be dominated by cooperators or defectors (analogous to mutualists and parasites in this model) depending on the initial conditions (Doebeli et al. 2004). Therefore, stochastic processes may be key in navigating between parasitic and mutualistic equilibria and are prime targets for future analyses.

We have also assumed no evolution on the part of the host. Host evolution is likely a key factor in the evolution of defensive symbioses and could interact with symbiont evolution in several ways. Hosts may evolve to minimize the costs associated with hosting symbionts by evolving tolerance (Carval and Ferriere 2010), resulting in lower virulence effects and cost to the symbiont. Symbiotic defense of the host could also release a host from having to defend against enemies (Nomura et al. 2010). Additionally, diversity within the symbiotic community could maintain variation in host traits for defense or tolerance of symbionts (Heil et al. 2009). As we have shown that evolution of just two symbiont traits can produce counterintuitive results, the outcomes of co-evolution between hosts and defensive symbionts are likely to be particularly hard to predict. Including host evolution may further elucidate the mechanisms by which benign symbioses are maintained in symbiotic communities.

### Conclusion

We have put forth a model in which symbionts can affect their hosts through their effect on host mortality, via virulence, and by defending the host against other symbionts. We have demonstrated that evolved defense largely leads to less damaging pathogens and preserves mutualisms under co-infection of the host by multiple symbionts. However, the evolution of defense can also lead to more deleterious pathogens and disrupt mutualisms if the costs and benefits affect symbiont reproduction and mortality differently. Thus, while defensive traits in symbionts may be key to maintaining benign symbioses, understanding how defensive traits affect the host and symbiont is vital to understanding the evolution of the host-symbiont relationship.

## References

Abrams, P. A., H. Matsuda, and Y. Harada. 1993. Evolutionarily unstable fitness maxima and stable fitness minima of continuous traits. Evol. Ecol. 7:465–487.

Afkhami, M. E., J. A. Rudgers, and J. J. Stachowicz. 2014. Multiple mutualist effects: Conflict and synergy in multispecies mutualisms. Ecology 95:833–844.

Alizon, S., A. Hurford, N. Mideo, and M. Van Baalen. 2009. Virulence evolution and the trade-off hypothesis: History, current state of affairs and the future. J. Evol. Biol. 22:245–259.

Anderson, R. M., and R. M. May. 1978. Regulation and stability of host-parasite population interactions: I. Regulatory processes. J. Anim. Ecol. 47:219–247.

Arnold, A. E., L. C. Mejia, D. Kyllo, E. I. Rojas, Z. Maynard, N. Robbins, and E. A. Herre. 2003. Fungal endophytes limit pathogen damage in a tropical tree. Proc. Natl. Acad. Sci. USA 100:15649–15654.

Balmer, O., S. C. Stearns, A. Schötzau, and R. Brun. 2009. Intraspecific competition between co-infecting parasite strains enhances host survival in african trypanosomes. Ecology 90:3367–3378.

Barrett, L. G., T. Bell, G. Dwyer, and J. Bergelson. 2011. Cheating, trade-offs and the evolution of aggressiveness in a natural pathogen population. Ecol. Lett. 14:1149–1157.

Betts, A., C. Rafaluk, and K. C. King. 2016. Host and parasite evolution in a tangled bank. Trends Parasitol. 32:863–873.

Bordenstein, S. R., and K. R. Theis. 2015. Host biology in light of the microbiome: Ten principles of holobionts and hologenomes. PLoS Biol. 13:e1002226.

Bremermann, H. J., and J. Pickering. 1983. A game-theoretical model of parasite virulence. J. Theor. Biol. 100:411–426.

Broyden, C. G. 1965. A class of methods for solving nonlinear simultaneous equations. Mathematics of computation 19:577–593.

Buckling, A., and M. Brockhurst. 2008. Kin selection and the evolution of virulence. Heredity 100:484–488.

Carroll, G. 1988. Fungal endophytes in stems and leaves: From latent pathogen to mutualistic symbiont. Ecology 69:2–9.

Carval, D., and R. Ferriere. 2010. A unified model for the coevolution of resistance, tolerance, and virulence. Evolution 64:2988–3009.

Choisy, M., and J. C. de Roode. 2010. Mixed infections and the evolution of virulence: Effects of resource competition, parasite plasticity, and impaired host immunity. Am. Nat. 175:>E105–118.

Clay, K., and C. Schardl. 2002. Evolutionary origins and ecological consequences of endophyte symbiosis with grasses. Am. Nat. 160:S99–S127.

Debarre, F., S. L. Nuismer, and M. Doebeli. 2014. Multidimensional (co)evolutionary stability. Am. Nat. 184:158–171.

Doebeli, M., C. Hauert, and T. Killingback. 2004. The evolutionary origin of cooperators and defectors. Science 306:859–862.

Ebert, D., and K. L. Mangin. 1997. The influence of host demography on the evolution of virulence of a microsporidian gut parasite. Evolution 51:1828–1837.

Ewald, P. W. 1987. Transmission modes and evolution of the parasitism-mutualism continuum. Ann. N. Y. Acad. Sci. 503:295–306.

Fenton, A., K. N. Johnson, J. C. Brownlie, and G. D. Hurst. 2011. Solving the wolbachia paradox: Modeling the tripartite interaction between host, wolbachia, and a natural enemy. Am. Nat. 178:333–342.

Ford, S. A., D. Kao, D. Williams, and K. C. King. 2016. Microbe-mediated host defence drives the evolution of reduced pathogen virulence. Nat. Comm. 7:13430.

Ford, S. A., and K. C. King. 2016. Harnessing the power of defensive microbes: Evolutionary implications in nature and disease control. PLoS Path. 12:e1005465.

Frank, S. A. 1996. Models of parasite virulence. Q. Rev. Biol.:37–78.

Friesen, M. L., and A. Mathias. 2010. Mixed infections may promote diversification of mutualistic symbionts: Why are there ineffective rhizobia? J. Evol. Biol. 23:323–334.

Gerardo, N. M., and B. J. Parker. 2014. Mechanisms of symbiont-conferred protection against natural enemies: An ecological and evolutionary framework. Curr Opin Insect Sci 4:8–14.

Heil, M., M. Gonzalez-Teuber, L. W. Clement, S. Kautz, M. Verhaagh, and J. C. Bueno. 2009. Divergent investment strategies of acacia myrmecophytes and the coexistence of mutualists and exploiters. Proc. Natl. Acad. Sci. USA 106:18091–18096.

Heithaus, E. R., D. C. Culver, and A. J. Beattie. 1980. Models of some ant-plant mutualisms. Am. Nat. 116:347–361.

Jaenike, J., R. Unckless, S. N. Cockburn, L. M. Boelio, and S. J. Perlman. 2010. Adaptation via symbiosis: Recent spread of a drosophila defensive symbiont. Science 329:212–215.

Johnson, N., J. H. Graham, and F. Smith. 1997. Functioning of mycorrhizal associations along the mutualism–parasitism continuum. New Phytol. 135:575–585.

Jones, E. O., A. White, and M. Boots. 2007. Interference and the persistence of vertically transmitted parasites. J. Theor. Biol. 246:10–17.

Jones, E. O., A. White, and M. Boots. 2011. The evolution of host protection by vertically transmitted parasites. Proc. R. Soc. Lond., Ser. B: Biol. Sci. 278:863–870.

Kerr, B., M. A. Riley, M. W. Feldman, and B. J. Bohannan. 2002. Local dispersal promotes biodiversity in a real-life game of rock–paper–scissors. Nature 418:171–174.

Kirkup, B. C., and M. A. Riley. 2004. Antibiotic-mediated antagonism leads to a bacterial game of rock–paper–scissors in vivo. Nature 428:412–414.

Koskella, B., T. Giraud, and M. Hood. 2006. Pathogen relatedness affects the prevalence of within-host competition. Am. Nat. 168:121–126.

Kwiatkowski, M., and C. Vorburger. 2012. Modeling the ecology of symbiont-mediated protection against parasites. Am. Nat. 179:595–605.

Lenski, R. E., and R. M. May. 1994. The evolution of virulence in parasites and pathogens: Reconciliation between two competing hypotheses. J. Theor. Biol. 169:253–265.

Lipsitch, M., S. Siller, and M. A. Nowak. 1996. The evolution of virulence in pathogens with vertical and horizontal transmission. Evolution:1729–1741.

Lively, C. M., K. Clay, M. J. Wade, and C. Fuqua. 2005. Competitive co-existence of vertically and horizontally transmitted parasites. Evol. Ecol. Res. 7:1183–1190.

Madden, D., and T. P. Young. 1992. Symbiotic ants as an alternative defense against giraffe herbivory in spinescent acacia drepanolobium. Oecologia 91:235–238.

May, G. 2016. Here come the commensals. Am. J. Bot. 103:1709–1711.

May, G., and P. Nelson. 2014. Defensive mutualisms: Do microbial interactions within hosts drive the evolution of defensive traits? Funct. Ecol. 28:356–363.

Mosquera, J., and F. R. Adler. 1998. Evolution of virulence: A unified framework for coinfection and superinfection. J. Theor. Biol. 195:293–313.

Nahum, J. R., B. N. Harding, and B. Kerr. 2011. Evolution of restraint in a structured rock–paper–scissors community. Proc. Natl. Acad. Sci. USA 108:10831–10838.

Nelson, P., and G. May. 2017. Co-evolution between mutualists and parasites in symbiotic communities may lead to the evolution of lower virulence. Am. Nat. 190:803–817.

Neuhauser, C., and J. E. Fargione. 2004. A mutualism–parasitism continuum model and its application to plant–mycorrhizae interactions. Ecol. Model. 177:337–352.

Nomura, M., A. Hatada, and T. Itioka. 2010. Correlation between the leaf turnover rate and anti-herbivore defence strategy (balance between ant and non-ant defences) amongst ten species of macaranga (euphorbiaceae). Plant Ecol. 212:143–155.

Oliver, K. M., J. Campos, N. A. Moran, and M. S. Hunter. 2008. Population dynamics of defensive symbionts in aphids. Proc. R. Soc. Lond., Ser. B: Biol. Sci. 275:293–299.

Orivel, J., P. J. Male, J. Lauth, O. Roux, F. Petitclerc, A. Dejean, and C. Leroy. 2017. Trade-offs in an antplant-fungus mutualism. Proc Biol Sci 284.

Prado, F., and B. Kerr. 2008. The evolution of restraint in bacterial biofilms under nontransitive competition. Evolution 62:538–548.

Press, W. H., S. A. Teukolsky, W. T. Vetterling, and B. P. Flannery. 2007, Numerical recipes 3rd edition: The art of scientific computing, Cambridge university press.

Saikkonen, K. 2007. Forest structure and fungal endophytes. Fungal Biology Reviews 21:67–74.

Sorrell, I., A. White, A. B. Pedersen, R. S. Hails, and M. Boots. 2009. The evolution of covert, silent infection as a parasite strategy. Proc. R. Soc. Lond., Ser. B: Biol. Sci. 276:2217–2226.

van Baalen, M., and M. W. Sabelis. 1995. The dynamics of multiple infection and the evolution of virulence. Am. Nat. 146:881–910.

Verbruggen, E., C. El Mouden, J. Jansa, G. Akkermans, H. Bucking, S. A. West, and E. T. Kiers. 2012. Spatial structure and interspecific cooperation: Theory and an empirical test using the mycorrhizal mutualism. Am. Nat. 179:E133–146.

Vittecoq, M., C. Djiéto-Lordon, D. McKey, and R. Blatrix. 2012. Range expansion induces variation in a behavioural trait in an ant–plant mutualism. Acta Oecol. 38:84–88.

West, S. A., E. T. Kiers, E. L. Simms, and R. F. Denison. 2002. Sanctions and mutualism stability: Why do rhizobia fix nitrogen? Proc. R. Soc. Lond., Ser. B: Biol. Sci. 269:685–694.

